# Parallel processing of olfactory and mechanosensory information in the honeybee antennal lobe

**DOI:** 10.1101/2021.05.11.443598

**Authors:** Ettore Tiraboschi, Luana Leonardelli, Gianluca Segata, Elisa Rigosi, Albrecht Haase

**Affiliations:** Department of Physics, University of Trento, Trento, 38122, Italy; Center for Mind/Brain Sciences (CIMeC), University of Trento, Rovereto, 38068, Italy; Department of Electrical, Electronic, and Information Engineering, University of Bologna, Bologna, 40126, Italy; Department of Biology, Lund University, Lund, 22362, Sweden

**Keywords:** Mechanosensing, honeybee, Apis mellifera, antennal lobe, mechanosensory neurons, calcium imaging

## Abstract

We report that airflow produces a complex activation pattern in the antennal lobes of the honeybee *Apis mellifera*. Glomerular response maps provide a stereotypical code for the intensity and the dynamics of mechanical stimuli that is superimposed on the olfactory code. We show responses to modulated stimuli suggesting that this combinatorial code could provide information about the intensity, direction, and dynamics of the airflow during flight and waggle dance communication.

## Introduction

The antennal lobe (AL) is the insect neuropil associated with the encoding of odour information received via the antennas^1^. Odour molecules bind to the olfactory receptors in the dendrites of olfactory receptor neurons (ORNs). In the honeybee *Apis mellifera*, each class of ORNs projects into a single of the 160 nodes of the AL network, called glomeruli. Those are interlinked by local neurons and project a stereotypical activation pattern, coding for odour identity and concentration^<u>2</u>^ to higher-order brain centres like the mushroom bodies (MBs) and the lateral horns (LHs)^1^. Besides the odour receptors located in the sensilla, hair-like structures exposed to ambient airflow, the antenna houses further receptors involved in sensing humidity, temperature^3^, and mechanosensory stimuli^4^. The latter are known to be detected in the pedicel of the antenna by Johnston’s organ, whose neurons project into the dorsal lobe^5^. A limited mechanosensitivity in the antennal lobe of moths^6–11^ was already reported. To clarify the involvement of the antennal lobe in mechanosensation, we systematically investigated glomerular responses during exposure to different air currents with or without additional odour stimuli using two-photon calcium imaging.

## Results

Flow rates were chosen to simulate what a bee would typically experience during flight (high flux, HF) and what wing beating during the waggle dance would produce (low flux, LF)^12^.

Exposed to repeated airflow stimuli (Fig. 1a), we observed in most of the imaged glomeruli clear and consistent responses (Fig. 1b, c, 5a, Supplementary Video S1). This confirms a hypothesis of Tuckman *et al*.^11^ that the glomerular mechanosensory response might be broader than that to single odours. But beyond previously reported activation^11^., we also found strong inhibition in several glomeruli (Fig. 1c, e). This suggests that the mechanosensitivity of receptor neurons is non-uniform and that probably the same inhibitory local neurons involved in odour coding generate these combinatorial patterns encoding airflow stimuli.

**Fig. 1.**
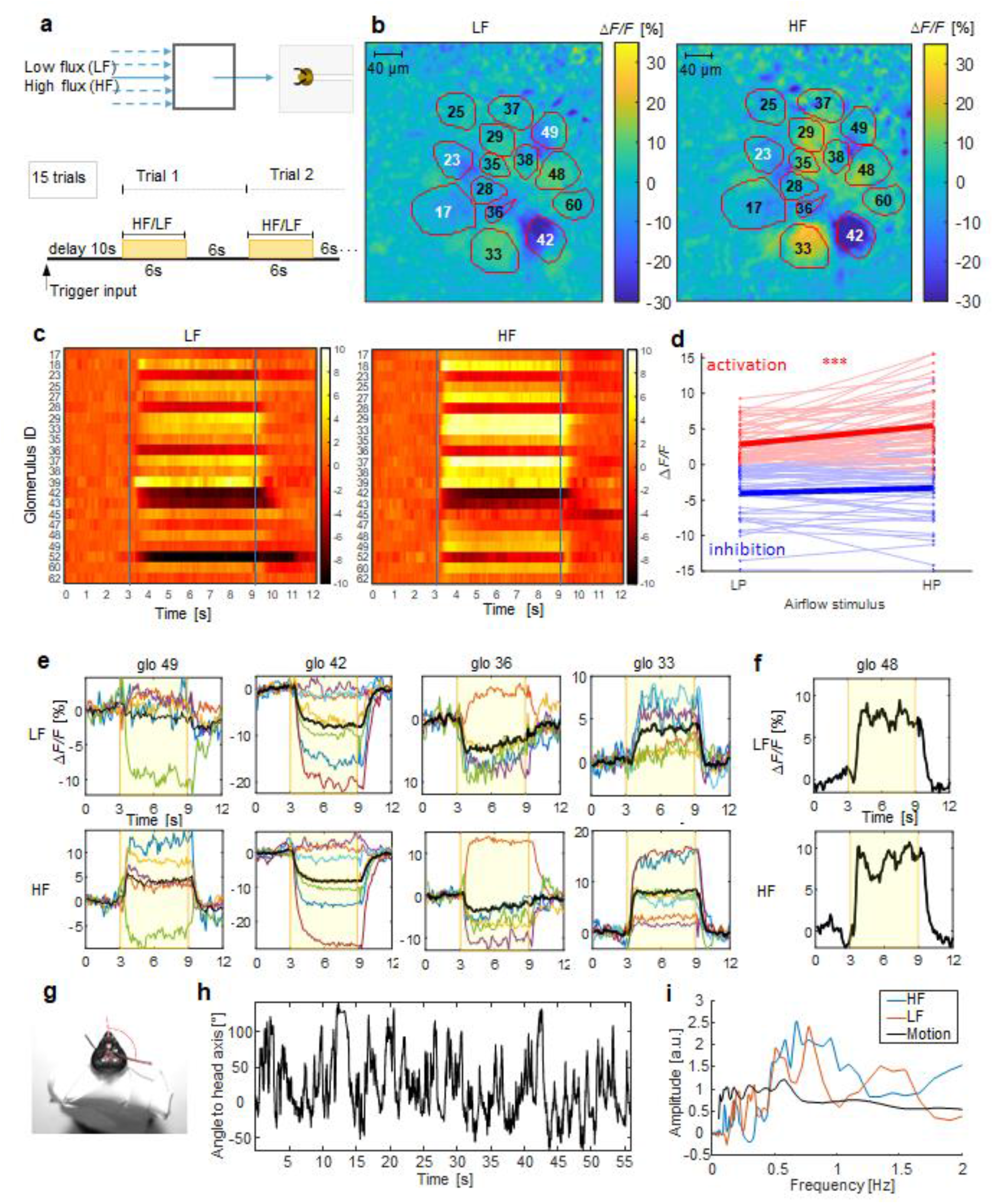
Response patterns to airflow stimuli. **a**, Setup scheme and stimulus protocol. Stimuli start after 10 s of background acquisition, lasting 6 s (yellow area), inter-trial distance 6 s. **b**, Example for the relative fluorescence change (Δ*F*/*F* [%]) in the imaging plane across the AL, outlines and labels show the identified glomeruli. **c**, Heatmaps show subject-averaged (*N*=7) responses of all imaged glomerulus to low flux (LF) and high flux (HF) delivered after 3 s. **d**, Change of the glomerular activation between LF and HF, activated glomeruli (red dots) increase responses significantly (paired *t*-test: *t*(56) = −5.51, *p* = 10-7), inhibited glomeruli (blue dots) don’t (*t*(49) = −1.65, *p* = 0.11) **e**, Temporal response curves of 4 selected glomeruli to low flux (LF) and high flux (HF) airflow. Coloured curves show single subject responses, averaged over 15 repetitions. Black curve is the subject-averaged response. The yellow background marks the stimulus interval. **f**, Example of glomeruli showing an oscillatory modulation of the activity signal, which was well conserved across the 15 trials. **g**, Bee mounted with the head and the antennas free to move for high-speed antenna motion imaging, current angle of the right flagellum is marked in red. **h**, Example for an antenna tracking curve during 1 min of recording. **i**, Averaged spectra of the oscillatory activity in (f) (red LF, blue HF) and spectrum of the antenna motion in (h).

To test the stereotypy of this code, the individual glomeruli were identified via the AL atlas^13^ and the experiment was repeated in 7 subjects. Results show that the response patterns are highly preserved across individuals (Fig. 1e, Extended Data Fig.1a, b). Comparing the distributions of glomerular responses between the stimuli and to the pre-stimulus activity, we found statistically significant differences between all of them (a PCA of the response distributions and statistical results are shown in Extended Data Fig. 2).

**Fig. 2.**
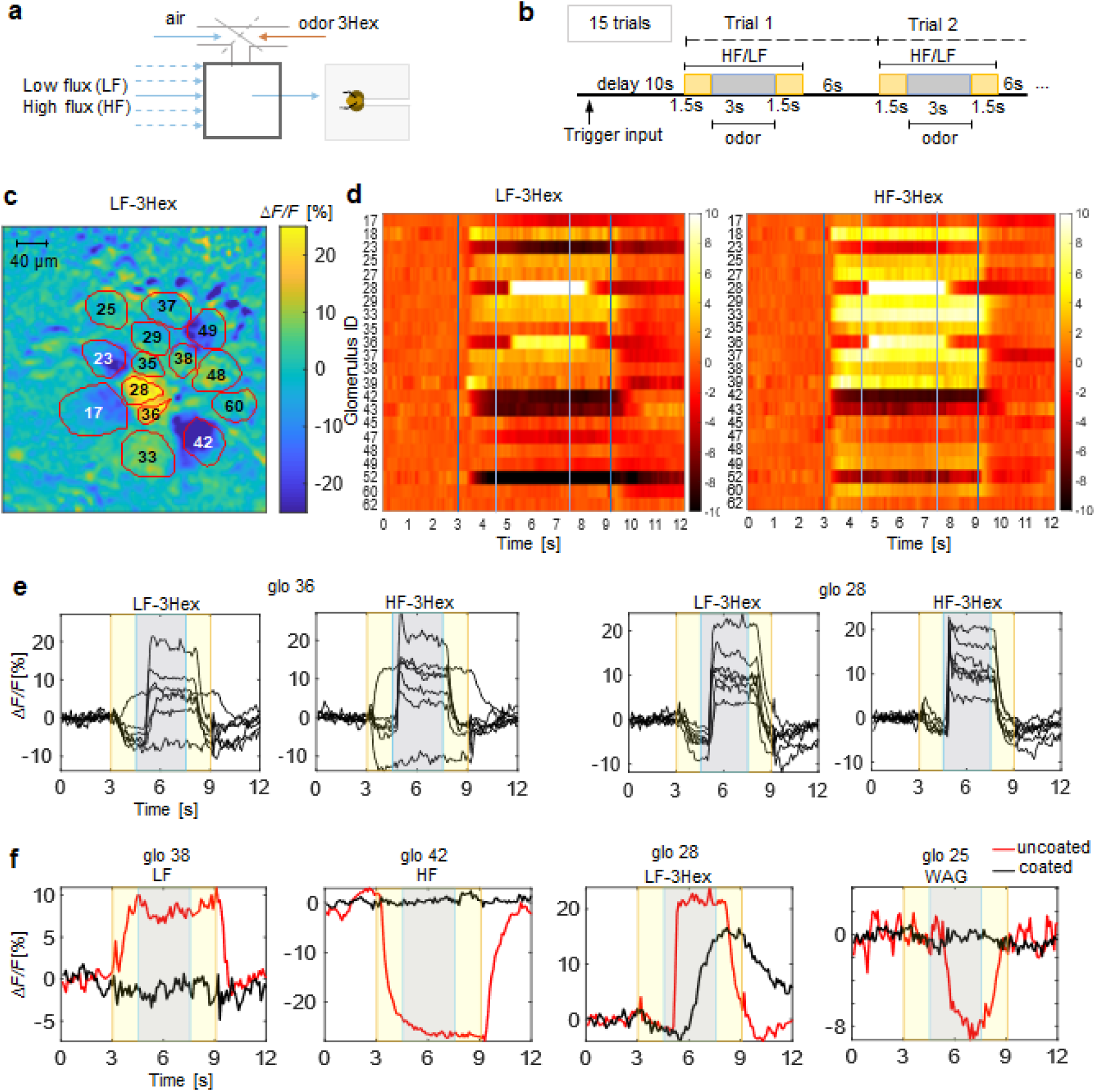
Response patterns to mechanical and chemical stimuli. **a**, Scheme of the setup where either clean air or 3-hexanol (3Hex) is injected into the carrier flux. **b**, Scheme of the stimulation protocol: To the airflow stimulus starting after 10 s (yellow area), the 3-Hex odour stimulus is added after another 1.5 s lasting 3 s (grey area), interstimulus interval 6 s. **c**, Relative fluorescence change in the imaging plane during the low air flux plus odour stimulus (LF-3Hex), outlines and labels show the identified glomeruli. **d**, Heatmaps show subject-averaged responses of all imaged glomerulus to low flux (LF) and high flux (HF) delivered after 3 s and air plus odour after 4.5 s **e**, Temporal response curves of the two glomeruli (28,36) that showed responses to the odour stimulus, yellow areas mark the air only periods, grey boxes the air plus odour periods. **f**, Temporal response curves of 4 selected glomeruli to odour, weak flow, high flow and waggling for bees with antennas coated with fluid silicon (black) and uncoated antennas (red).

We also observed that glomerular activation is proportional to the airflow intensity whereas glomerular inhibition is not (Fig. 1d). The glomerular response to the airflow rarely attenuates during the 6 s of exposure (Fig. 1e), in contrast to odour responses which usually decrease over time. This lack of habituation suggests that continuous monitoring of wind speed during flight could be based on this code.

A particular case of signal modulation is shown in Fig. 1f, where the glomerular response shows an oscillatory modulation, which is consistently reproduced during the 15 trials. Comparing this modulation with the angular motion of the flagellum obtained by high-speed video tracking (Fig. 1g, h), we found good agreement between the frequency spectra of neuronal activity and motion in the region that characterizes the oscillatory modulation (Fig.1i). The reorientation of the flagella changes the direction under which the airflow hits the sensilla. This seems to produce a direction-dependent signal modulation. Bees might use this direction-sensitivity not only to detect odour gradients, as observed in cockroaches^14^, but also to sample the wind direction during flight.

We then studied the responses elicited by a superposition of mechanical and odour stimuli by injecting 3-Hexanol (3Hex) into the air stream (Fig. 2a, b), an odour that is known to excite glomeruli 28 and 36^15^. In this experiment, the odour stimulus lasted 3 s and was added after 1.5 s to the airflow stimulus, without changing the overall flux (Fig. 2c, d, Extended Data Fig. 2b, c). Both glomeruli, which are initially inhibited by the airflow, show a reversal of this inhibition into a strong activation (Fig.2c, d, e, Supplementary Video 2). This shows that mechanosensory responses do not necessarily reduce the odour signal contrast. At realistic airflow velocities, chemosensation was found to be dominant, which can be expected since airflow is a rather continuous stimulus during flight, whereas olfactory stimuli are sparse, highly variant, and of great relevance and therefore require precedence in perception.

To verify the origin of both signals, we coated the flagellum with a thin layer of silicone, but leaving it free to move, and repeated the experiment. The observed mechanosensory response was now highly attenuated, whereas the odour response was as strong as before although with slower response dynamics (Fig. 2f). Silicone slows down the diffusion of the odour molecules toward the chemoreceptors and strongly impairs mechanoreceptors activation. This rules out that the mechanosensory signal in the glomeruli originates from Johnston’s organ, as the flagellum could move freely. We hypothesize that it is rather the motion of the sensilla which was damped by the silicone coating that mediates the mechanosensitivity.

Next, we tested whether the AL mechanosensation has the potential to play a role in the waggle dance communication, where dancer bees communicate angle and distance of a food source by wing beating and abdominal oscillations to dance followers^16^. Michelsen *et al*^.*<u>12</u>*^ reported an airflow elicited by the wing beating from 0.15 to 0.3 m/s modulated by abdominal oscillations at a frequency of 24-25 Hz. We reproduced a waggle-dance-like stimulus (WAG) by oscillating a winglet at 24 Hz in a laminar airflow of 0.25 m/s (Fig. 3a, b). Already the oscillating winglet by itself elicited a stereotypical response in most glomeruli, either activating or inhibiting them (Fig. 3c, d, Extended Data Fig. 1c, d, Supplementary Video 3). The airflow generated by the winglet is very weak (average speed 0-0.03 m/s) however being very turbulent, it may generate strong local gradients leading to a pulsed-like stimulation of vibrational movements of the sensilla. We then embedded the waggle stimulus in a laminar airflow, reproducing precisely the airflow felt by a bee that is following the dancer. The results clearly show that this modulation of the laminar airflow is effectively detected by the glomeruli (Fig. 3c, d). The waggle stimulus is more effective in a slow airflow and we observed different characteristics in the glomerular responses. We found glomeruli sensitive already to the waggle stimulus without an airstream (Fig. 3e, g, h), others were sensitive only to a combined waggling/airflow stimulus (Fig. 3f). Some were tuned to detect waggling in a weak flow but not in the strong flow (Fig. 3e) and others again were modulated either in a weak or strong flow (Fig. 3f, g). This rich repertoire of responses suggests a high dynamic range of the mechanoreception mechanism which would allow for coding of complex temporal patterns (Extended Data Fig. 3, 4 show the complete spatio-temporal response pattern to all stimuli in a representative bee).

**Fig.3.**
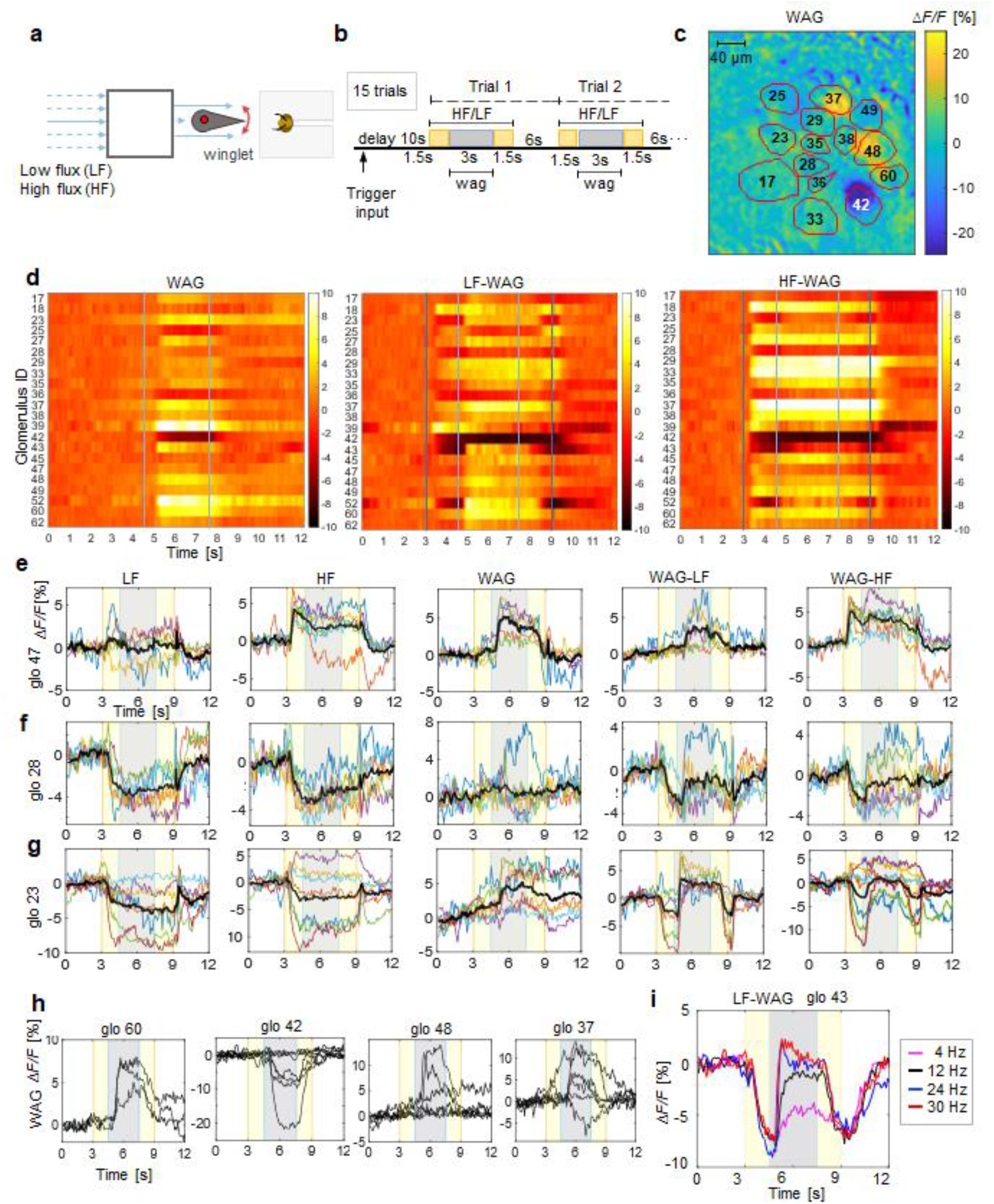
Response patterns to waggle motion. **a**, Stimulus generator scheme for laminar airflow and waggle-dance-like stimuli via an oscillating winglet. **b**, Stimulation protocol with laminar flow stimuli starting after 10 s of background acquisition, lasting 6 s (yellow area) and a waggle motion added to it after 1.5 s lasting 3 s (grey areas), inter-stimulus interval 6 s. **c**, Relative fluorescence change in the imaging plane during stimulus only by the waggle motion (WAG) without additional airflow, outlines and labels show the identified glomeruli. **d**, Heatmaps show the subject-averaged glomerular responses to the waggle only stimulus (WAG) and combined stimuli where waggling is added after 4.5 s to the low flux (LF-WAG) or the high flux (HF-WAG). **e**, Temporal response curves to single and combined stimuli of glomerulus 47 which is sensitive already to waggling only. **f**, Temporal response curves to single and combined stimuli of glomerulus 28 not sensitive to waggling only, where waggling stronger modulates the LF stimulus. **g**, Temporal response curves to single and combined stimuli of glomerulus 23 sensitive to waggling and where waggling stronger modulates the HF stimulus. **h**, Temporal response curves of 4 selected glomeruli to waggle motion only. **i**, Response of a selected glomerulus to a low flux stimulus with superimposed waggle motion at different frequencies.

**Fig. 4.**
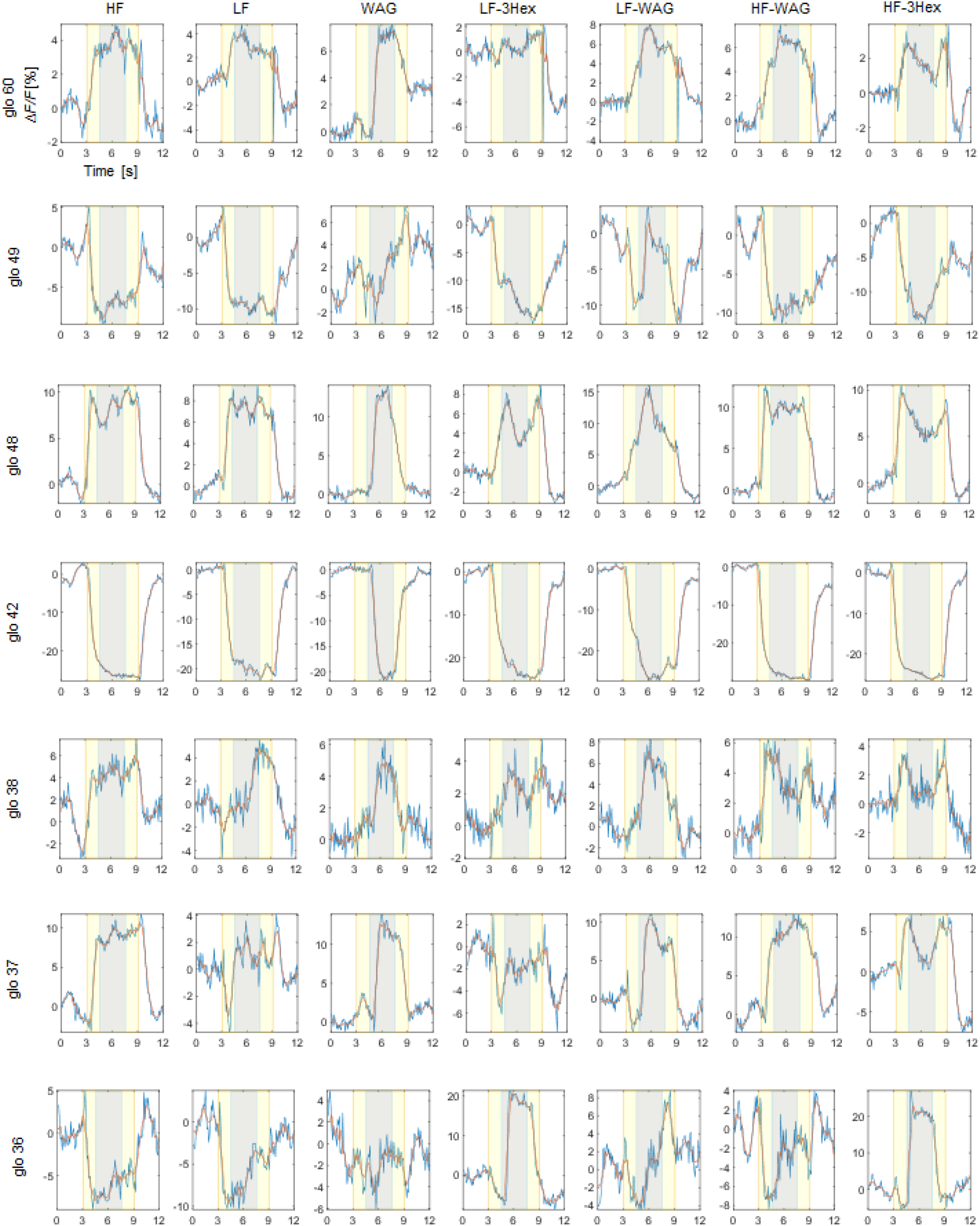
Complete set of stimuli of a representative bee (glo 36-glo 60). Rows show individual glomeruli, columns the different stimuli. Blue lines represent the response averaged over all 15 trials, the red line shows the low-pass filtered response. The yellow areas highlight the airstream stimulus period, the grey area the additional odour or waggle stimuli. The stimulus order is the same as provided during the experiment. The signal intensity expressed as Δ*F/F* [%].

We finally asked how the modulation of glomerular activity varied with the winglet oscillation frequency. We, therefore, repeated the experiment in a weak airflow and tested frequencies of 4, 12, 24, and 30 Hz. The results show that the strongest modulatory effect is observed already at 24 Hz with no further improvements at 30 Hz, whereas for lower frequencies the effect is proportionally reduced (Fig. 3i, Supplementary Video 4).

Since during waggle dance a bee produces oscillations also at higher frequencies at 200-400 Hz via wing beating, we tested the sensitivity of the AL also to these signals. We exposed the bees to comparable stimuli produced by a loudspeaker ramping frequencies from 40 up to 6000 Hz. We never observed a glomerular response to these signals.

## Discussion

In summary, these findings provide the first evidence of parallel coding of chemical and mechanical stimuli in the honeybee AL. So far studies have suggested Johnston’s organ as the major contributor to mechanosensation. We hypothesize that the glomerular code is likely contributing considerably to it. The observed response patterns show that mechanosensitive responses do not just amplify an odour signal but are, due to the broad response spectrum, their complex dynamics and, above all, their stereotypy, capable of encoding information relative to wind speed and direction. The tonic nature of the responses suggests that glomeruli record this information persistently but without reducing neither contrast nor the dynamic range of the chemical signals that are perceived in parallel. One type of stimuli to which the AL was found to be particularly sensitive, was periodic low-frequency modulations of the airflow, as they occur during the waggle dance communication. Interestingly, the response to oscillations reaches a maximum at 24 Hz, a frequency that was reported to provide the most efficient transfer of information during the waggle dance^12^. This happens when the follower bees are aligned within 30° to the dancer bee’s body axis. If instead the receiver bee is located laterally to the dancer, the oscillation frequency drops by one half to ca. 12 Hz and the information transfer was found to be less effective^12^. Our study supports this observation since at 12 Hz the modulatory effect on the activity was strongly reduced. On the other hand, it did not increase considerably at frequencies beyond 24 Hz. This potential involvement of the antennal lobe in waggle motion detection adds a further option by which higher brain centres might decipher the numerical information about the distance of a food source. Further studies could bring us closer to answering one of the most interesting questions in animal communication.

Overall, this study contributes to a new understanding of the olfactory system, as a network involved in the processing of a much broader spectrum of airborne information beyond odour identification^17^. The findings should also provide additional arguments for the importance of the honeybee as a neuroethological model for olfaction, as mechanosensitivity in olfactory neurons has recently also been discovered in mammals^18^. There are legitimate hopes that studies in a network of a few thousand neurons will contribute significantly to the understanding of the underlying mechanisms.

## Methods

### Specimen preparation for *in vivo* calcium imaging

Honeybees were prepared following a well-established protocol^19^. The bees were exposed to CO_2_ for 30 s. The immobilized bees were then fixed onto a custom-made imaging stage, using soft dental wax (Deiberit 502, Siladent). A small rectangular window was cut into the cuticula. The glands and trachea covering the AL were moved aside and fura2-dextran, a calcium-sensitive fluorescent dye (Thermo-Fisher Scientific) dissolved in distilled water was injected into the antenno-cerebralis tracts, postero-lateral to the α-lobe using a pulled glass capillary. After the injection, the cuticula was fixed in its original position using n-eicosane (Sigma Aldrich). The bees were stored in a dark, cool, and humid place for 15 - 20 h to let the dye diffuse into the AL.

Just before the imaging session, antennas were blocked with a drop of n-eicosane on the pedicel leaving the flagellum free to move. The cuticular window, the trachea, and the glands were removed from the antennal lobe region. A silicone adhesive (Kwik-Sil, WPI) was used to cover the brain and a rectangular plastic foil was attached frontal to the window to separate the antennas from the immersion water for the objective lens.

### Two-photon microscope

The two-photon microscope (Ultima IV, Bruker) was illuminated by a Ti:Sa laser (Mai Tai Deep See HP, Spectra-Physics). The laser was tuned to 780 nm for fura-2 excitation. All images were acquired with a water immersion objective (10×, NA 0.3, Olympus). Fluorescence was collected in epi-configuration, selected by a dichroic mirror, filtered with a band-pass filter centred at 525 nm and with a 70 nm bandwidth (Chroma Technology), and detected by a photomultiplier tube (Hamamatsu Photonics). The laser power was limited to about 10 mW to reduce photodamage on the specimen, maintaining a good signal-to-noise (SNR) ratio.

### Mechanosensory and odour stimulation

We built a custom device for delivering controlled odour and mechanosensory stimuli. Pure air from a pressure-controlled source passed a charcoal filter and was then humidified by a water flask. The airflow is switched with two solenoid valves in a serial configuration. The first valve opens and closes the airstream. When closed, the airstream is diverted into an exhaust channel to prevent pressure from building up in the system, which creates a rectangular stimulus profile without an initial spike after opening the valve. The second valve determines the flow rate by switching between a large or narrow duct. The airstream speed can be varied between 1.8 m/s (HF) and 0.25 m/s (LF) via a mechanical airflow meter (ANALYT-MTC) and is measured at the position of the antennas with a thermo-anemometer (testo 405i, Testo). Upstream there is a 3-way valve (LHDA0531115, The Lee Company) adding either the oil-immersed odour or pure air to the carrier stream, such that the overall airflow remains constant during the entire stimulation protocol (Fig. 2a). The airstream is aimed at the bee’s head via a steel tube of 15 cm length and 10 mm cross-section. centred in front of the steel tube is a vertical winglet (10×10 mm, L×H) which can oscillate laterally driven by a DC motor to produce a waggle stimulus. The winglet is coated with aluminium foil and grounded to earth to prevent electrostatic charges in the airstream. The distance between the winglet tip and the head of the bee is about 15 mm. The solenoid valves and the DC motor are controlled with an Arduino Uno board (Arduino) through custom software. Sound stimuli were generated using the Arduino Uno board, an audio amplifier board module (HiLetgo TDA2822M), and a speaker of 28 mm diameter, 8 Ω, and 2 W placed 15 mm frontally to the bee. Stimulation protocols were generated through MATLAB (R2019b, MathWorks) scripts and delivered to the Arduino board through a PCIe-6321 multifunction board (National Instruments). A recording session started with 10 s of background signal acquisition followed by alternating different types of stimuli in a pseudorandom order up to 15 trials per stimulus. The duration of the main mechanical stimulus was 6 sec, during which an airstream of different intensities, LF (0.25 m/s) HF (1.8 m/s) and no-air (0 m/s), was delivered. In the middle of this time window, a secondary stimulus of 3 s could be added (waggling WAG or odour 3Hex). The main stimulus period is followed by an interstimulus interval of 6 s. For the odour stimulus, we used 3-hexanol (W335118, Sigma-Aldrich), diluted 1:25 in mineral oil. Only the head of the bee is exposed to air/odour stimuli as the body is enclosed in the mounting stage to minimize mechanosensory stimulation of the insect body.

### Image acquisition

The image acquisition was synchronized to the stimulus protocol at a frame rate of 10.083 fps. The image of 128 × 128 pixels with a digital zoom factor of 3.8 covers a field of view of 280 µm. The fluorescence intensity was recorded with a depth of 13 bits. In addition to the functional images, a *z*-stack of the antennal lobe was acquired at a spatial resolution of 512×512 pixels and a layer interval of 2 µm to perform the morphological identification of glomeruli.

### Image analysis

A total of 7 bees were recorded and analyzed. Data post-processing and analysis were performed employing custom scripts in MATLAB. In each bee, the recorded glomeruli were identified using the AL atlas^<u>13</u>^ and associated with regions of interest (ROI) over which the fluorescence signals were spatially averaged. From these raw data the relative change of fluorescence during the stimulus and expressed in %: Δ*F*/*F*= −[*F*(t) − *F*_b_]/*F*_b_.×100, where *F*_b_ is the average fluorescence signal in the 3 s pre-stimulus period. This is a measure for the neuronal firing rate in each glomeruli^<u>20</u>^. Finally, for each stimulus, Δ*F*/*F* was averaged over the 15 trials to obtain the mean response for each glomerulus to a stimulus.

To identify glomeruli with the highest variance during the stimuli, a PCA was performed on the pixels as variables with frames as observations (Extended Data Fig. 5)

**Fig. 5.**
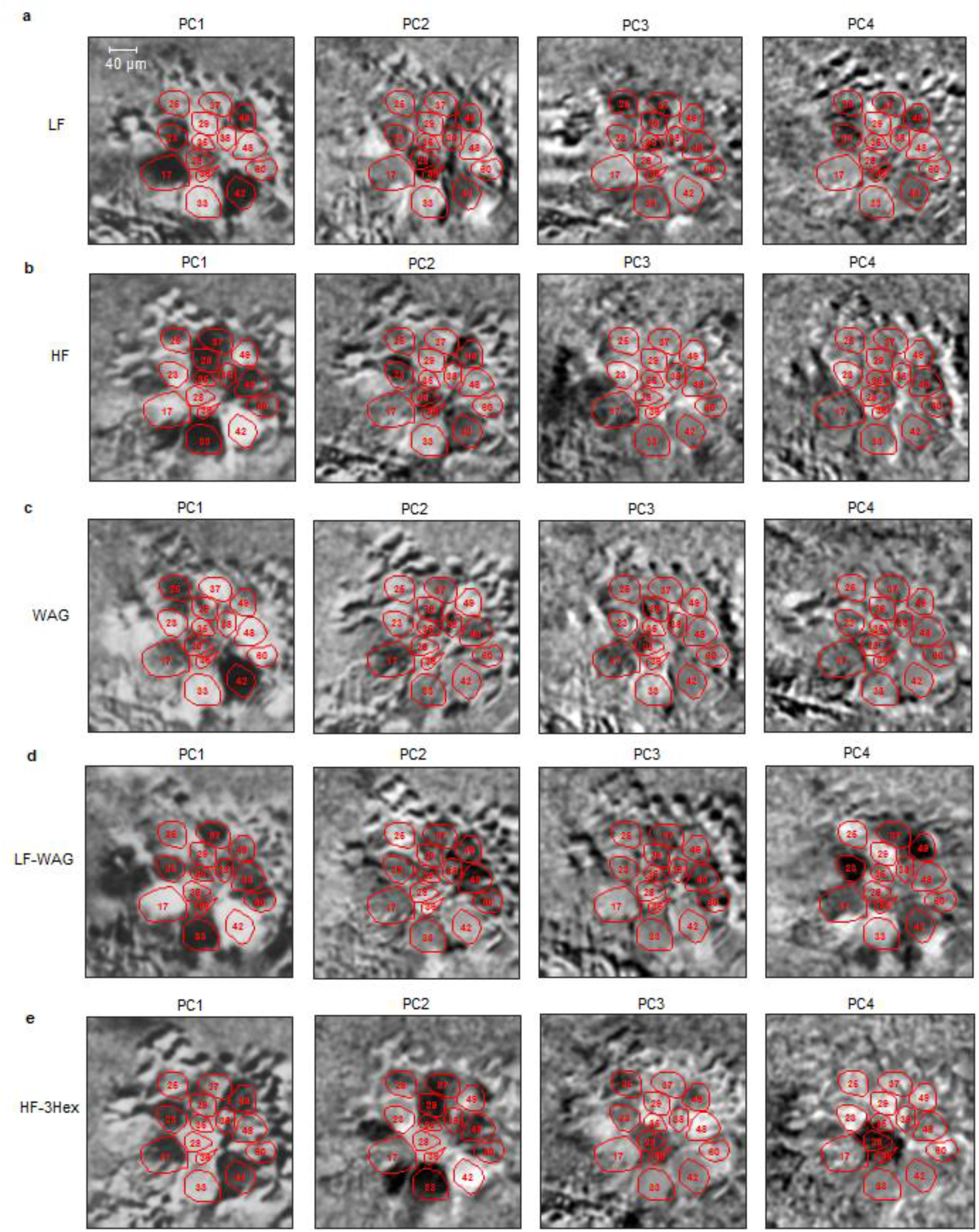
Principal components highlight glomeruli with the greatest signal variance during a stimulus. The frames of a stimulus period were averaged over trials, normalized, and converted into vectors. A PCA was then performed with pixels as variables and time frames as observations. The first PC is the variance-maximising projection of stimulus-related signals, spontaneous activity, and sample movements^21^. The strongest glomerular responses show high eigenvalues in the first principal component. Signals in the periphery are due to highly active neuronal somata and sample motion. The maps evidence the broad involvements of several glomeruli to stimuli encoding. **a**, Glomerular pattern elicited by LF stimulation. **b**, Glomerular pattern elicited by HF stimulation. **c**, Glomerular pattern elicited by waggling stimulation. **d**, Glomerular pattern elicited by LF airstream modulated by waggling. **e**, Glomerular pattern elicited by the odour 3-Hexanol.

### Statistical analysis of the stereotypy

A principal component analysis was performed on the full dataset using as features the averaged glomerular response in the first 1.5 s of each stimulus for the LF *vs*. HF *vs*. no-air comparison whereas for the LF *vs*. LF-3Hex the average over the last 2 s of the odour stimulus was used. The glomerular responses for no-air were computed averaging over 1.5 s before the stimulus. Every single recording corresponds to an observation. Statistical differences in the distribution of each group were evaluated using the statistical energy test. The multiple comparisons were corrected for familywise errors via the Bonferroni method.

### Antenna tracking

The antenna motion was recorded with a JVC GC-PX100BE Camcorder. A frame rate of 200 Hz turned out to be the best compromise between temporal and spatial resolution (640×360 pixels). Recordings were analyzed via custom python scripts. Images were background subtracted and binarized, and the antennas were identified via connected component analysis. Antenna images were then skeletonized, and the flagellum axis was obtained via the Hough transform. Its angle was measured against the head axis, which was obtained by polygonal fitting the head contour.

## Acknowledgement

We thank Paul Szyszka and Gustavo Borges Moreno e Mello for fruitful discussions and the University of Trento strategic project Brain Network Dynamics (BRANDY) for financial support. Elisa Rigosi acknowledges financial support from the Swedish Research Council for Sustainable Development (FORMAS 2018-01218).

## Author Contributions

E.T. designed the study, developed the methodology, acquired and analyzed the data. L.L. acquired the data and contributed to the data analysis. E. R. acquired the antenna motion data. G. S. analysed the antenna motion data. A.H. contributed to the data analysis. All authors contributed to the preparation of the manuscript.

## Declaration of Interests

The authors declare no competing interests.

## Supplementary Information

**Fig. 1.**
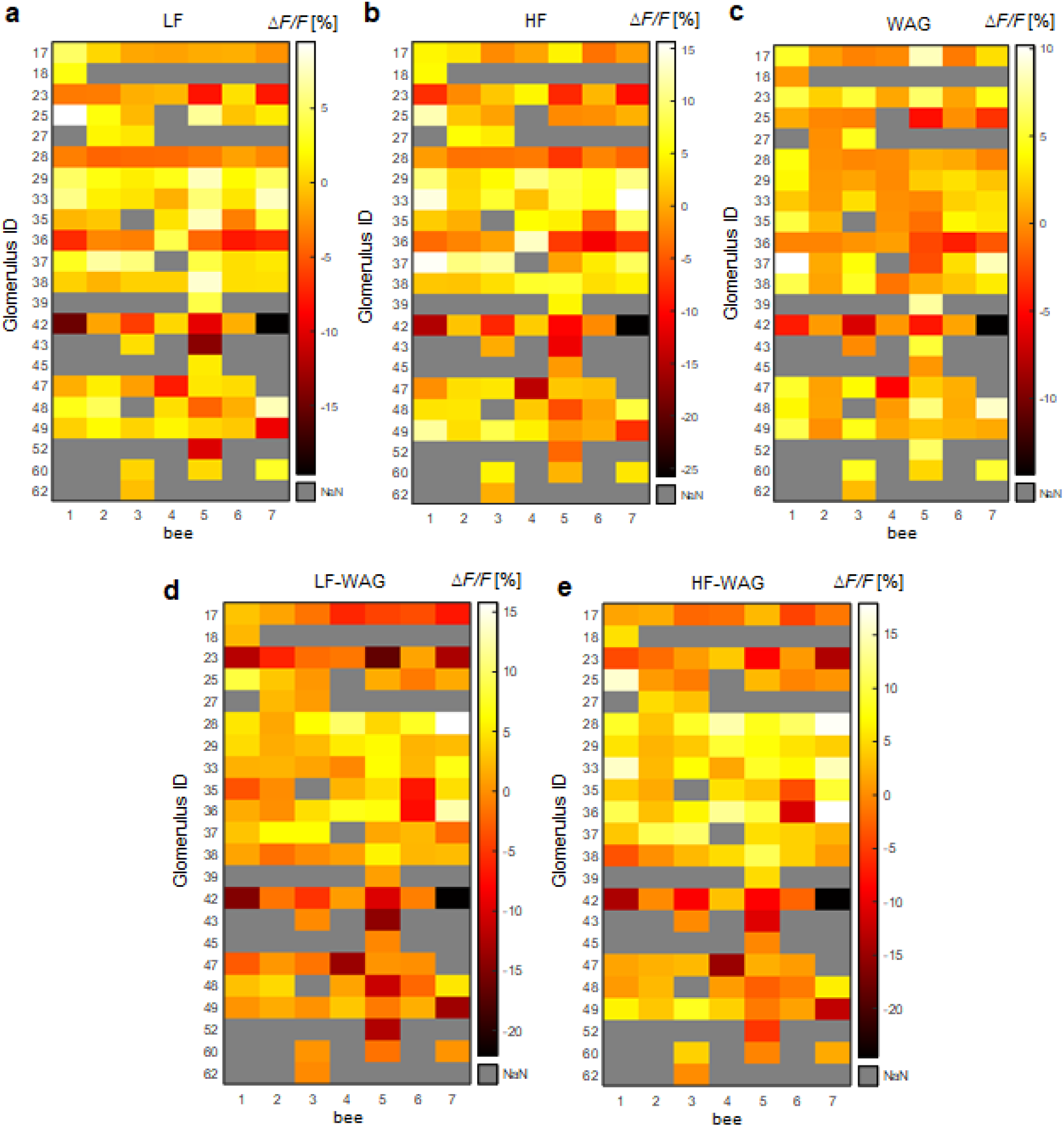
Maps of the glomerular responses to the different stimuli, for each recorded bee. Shown is the trial-averaged activity from 2 - 4 s after stimulus onset. Grey areas mark glomeruli that could not be recorded.

**Fig. 2.**
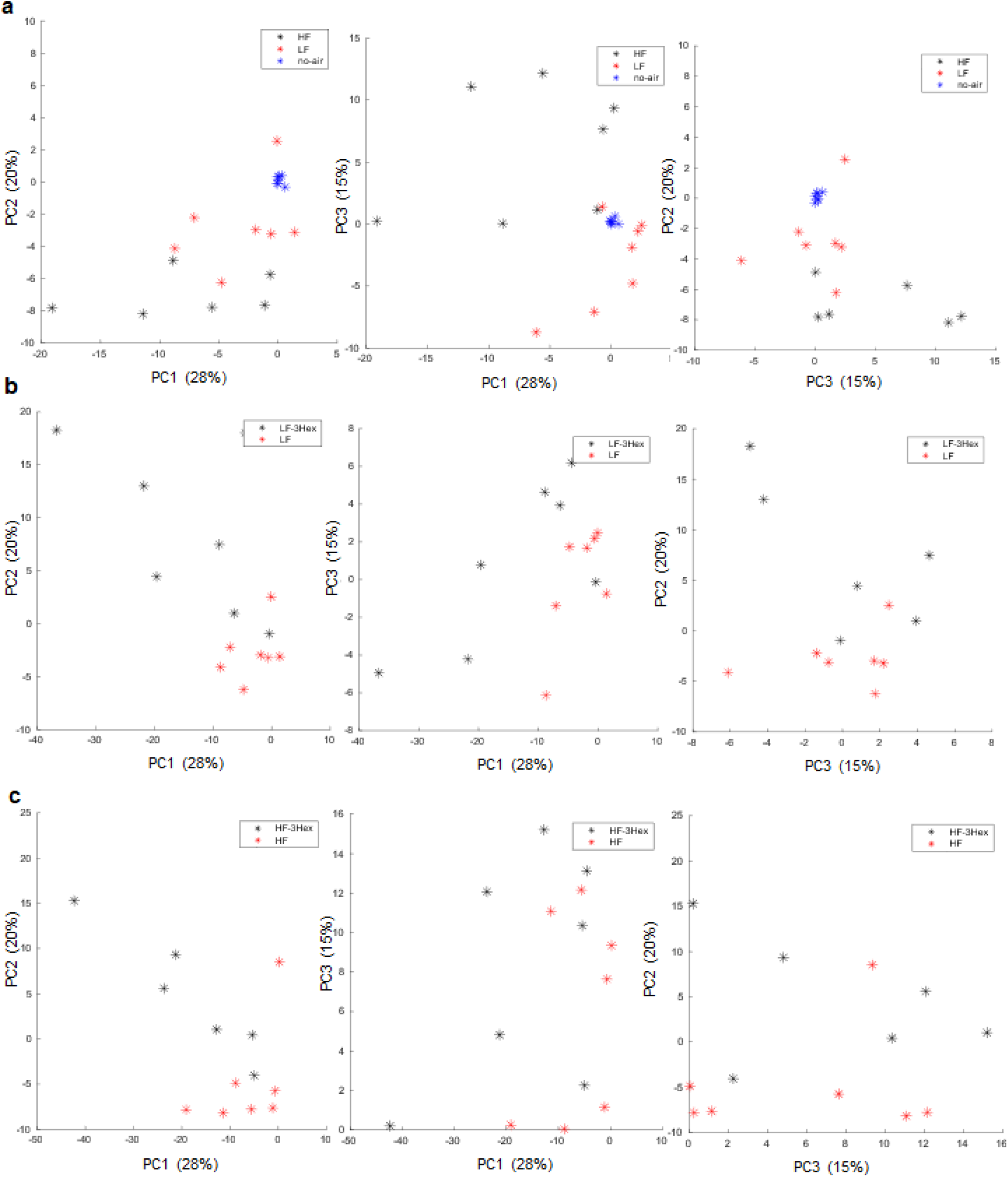
PCA of the glomerular response space. Each individual bee response to the stimuli is described in terms of the 3 first PCs. **a**, Comparisons between LF-HF-no-air stimuli. A statistical energy test^22^ gives HF *vs*. LF (𝜑(7) = 22.0, *p* = 0.022), no_air *vs*. LF (𝜑(7) = 18.9, *p* = 0.001), no-air *vs*. HF (𝜑(7) = 50.1, *p* = 0.001). **b**, Comparisons between LF and LF-3Hex (𝜑(7) = 51.3, *p* = 0.004). **c**, Comparisons between HF and HF-3Hex (𝜑(7) = 42.3, *p* = 0.037). All differences are significant including the Bonferroni correction for family-wise errors.

**Fig. 3.**
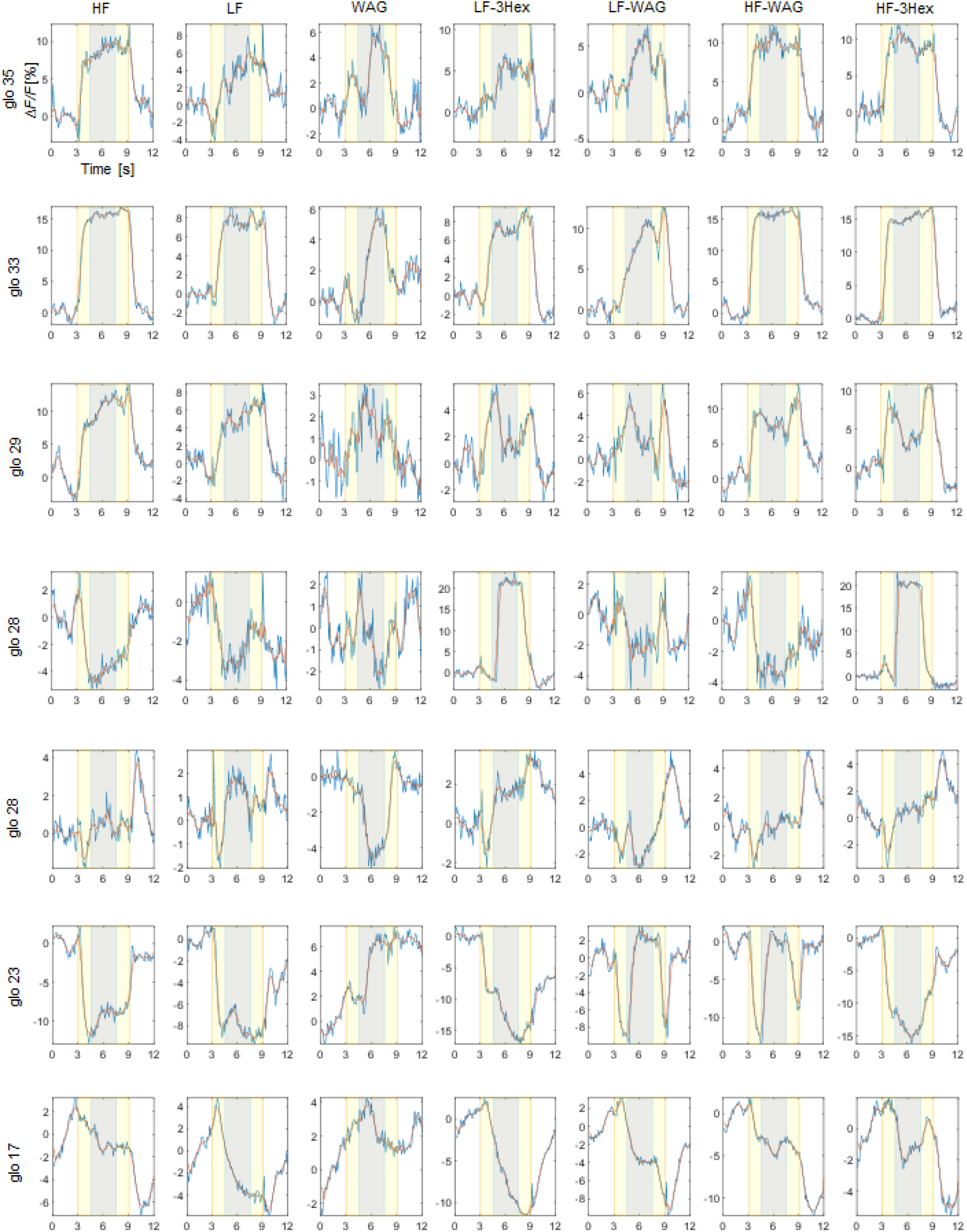
Complete set of stimuli of a representative bee (glo 17 - glo 35). Rows show individual glomeruli, columns the different stimuli. Blue lines represent the response averaged over all 15 trials, the red line shows the low-pass filtered response. The yellow areas highlight the airstream stimulus period, the grey area the additional odour or waggle stimuli. The stimulus order is the same as provided during the experiment. The signal intensity expressed as Δ*F/F* [%].

**Fig. 5.**
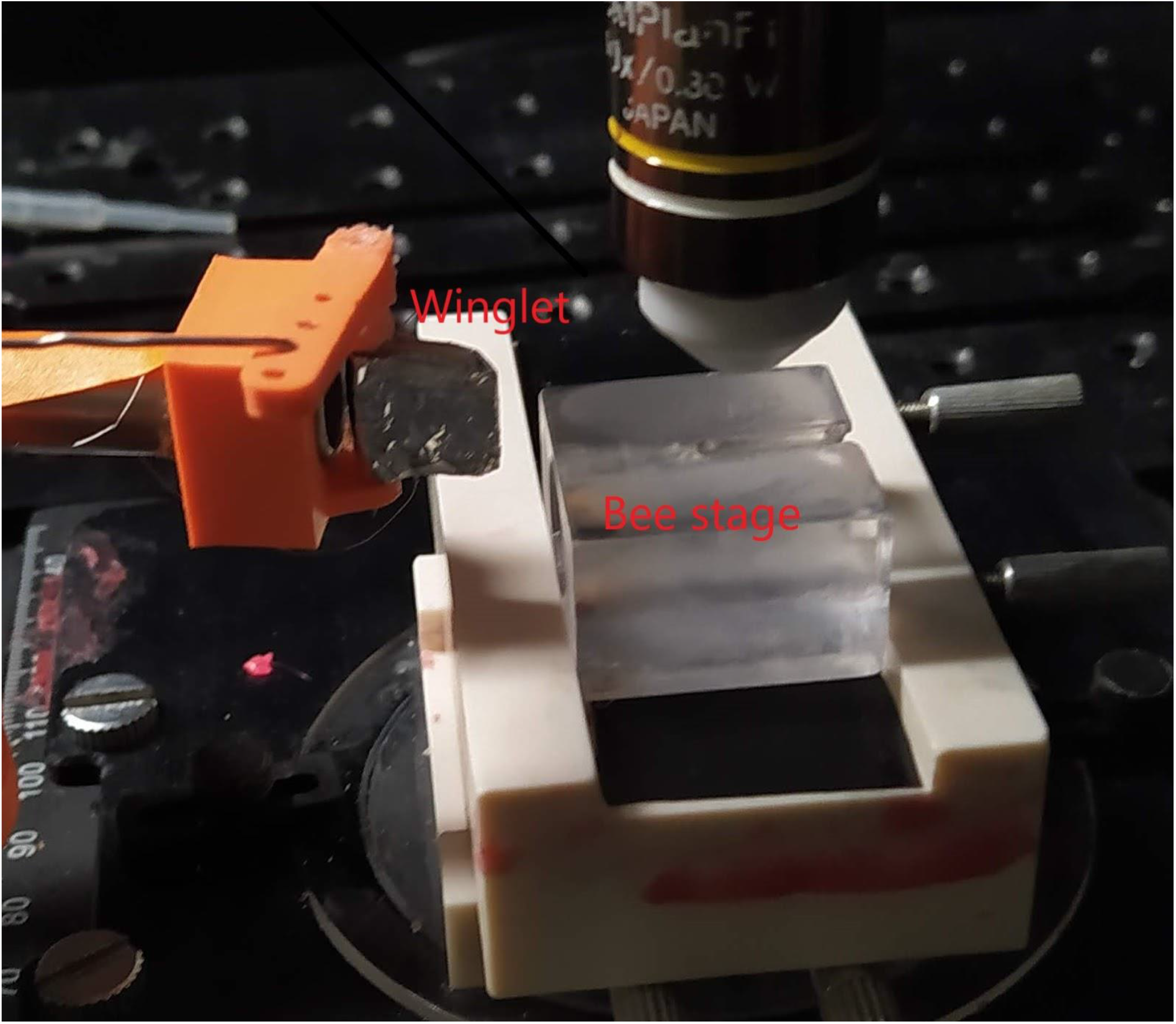
Stimuli generator. Bees are mounted on the stage facing the winglet. The airflow is directed through the steel tube straight toward the antennae. The winglet is activated by a motor whose speed is detected through a rotary encoder.

---

**Movie S1. Movie of the Glomerular responses to HF and LF stimuli**. Real-time trial-averaged response of a bee to the LF stimulus (left window) and the HF stimulus (right window), showing the higher response to the HF stimulus with respect to LF. The appearance of the red label indicates the stimulus period and type. Outlines and numbers indicate the glomeruli according to the bee atlas. The signal intensity is showing Δ*F/F* [%].

**Movie S2. Movie of the Glomerular responses to overlapping mechanical and olfactory stimuli**. Real-time trial-averaged glomerular responses to the LF-3Hex stimulus (left window) and the HF-3Hex stimulus (right window). The movie highlights the faster response to the odour (3-Hexanol) of glo28 compared to glo36 and shows the strong inhibition in glo17 and glo48 during odour stimulation in the LF but not in the HF airstream. This is an example of the complexity of the interaction between the mechano-and chemosensory encoding. The appearance of the red label indicates the stimulus period and type. The cyan label indicates the period of the odour delivery. Outlines and numbers indicate the glomeruli according to the bee atlas. The signal intensity is showing Δ*F/F* [%].

**Movie S3. Movie of the Glomerular responses to HF, LF and WAG stimulation**. Real-time trial-averaged responses of a bee to the LF-WAG stimulus (left window), the HF-WAG stimulus (right window) and to a simple WAG stimulus (central window). The central window shows the glomeruli which are modulated by simply oscillating the winglet at 24Hz. The appearance of the red label indicates the stimulus period and type. The cyan label indicates the waggling period of the winglet. Outlines and numbers indicate the glomeruli according to the bee atlas. The signal intensity is showing Δ*F/F* [%].

**Movie S4. Movie of the Glomerular responses to LF and WAG stimulation at different frequencies**. Real-time trial-averaged response recorded in the antennal lobe of a bee to the LF-WAG stimulus at 4 Hz (left window) and the LF-WAG stimulus at 24 Hz (right window). The appearance of the red label indicates the stimulus period and type. The cyan label indicates the waggling period. Outlines and numbers indicate the glomeruli according to the bee atlas. The signal intensity is showing Δ*F/F* [%].

## Notes

### Competing Interest Statement

The authors have declared no competing interest.

